# Temporal variability enhances the acquisition of stereotyped communication signals

**DOI:** 10.1101/2024.04.18.590102

**Authors:** Logan S. James, Olivia Ruge, Jon T. Sakata

## Abstract

Species-typical behaviors are organized into species-typical patterns, and deviations away from these patterns often diminish the strength of behavioral and sensory responses to such stimuli. In songbirds like the zebra finch, species-typical songs consist of acoustic elements (syllables) arranged into stereotyped (i.e., highly predictable) sequences with stereotyped timing. However, the degree to which deviations away from these stereotyped temporal patterns modulate the strength of vocal learning (i.e., the fidelity of vocal imitation) remains unknown. Here we tutored 123 juvenile zebra finches with stimuli that varied in the stereotypy of syllable sequencing and timing. In contrast to the prediction that deviations away from species-typical stereotypy would diminish vocal learning, deviations from sequence or timing stereotypy did not decrease how well juveniles imitated the acoustic structure of syllables. Moreover, presenting syllables in species-atypical sequences (i.e., randomized syllable sequences) enhanced vocal imitation in birds that were tutored later in development. This unexpected enhancement of birdsong learning by sequence variability resembles the effects of contextual diversity on speech acquisition and indicates that such variability can benefit learning even for very stereotyped behaviors.

**SIGNIFICANCE STATEMENT:** Little is known about the degree to which the temporal patterning of acoustic elements (“syllables”) modulates vocal learning and imitation. We tested the contribution of temporal variability in syllable sequencing and timing, hypothesizing that deviations away from species-typical stereotypy (predictability) in syllable sequence and timing patterns would diminish the strength of vocal learning. In contrast to our predictions, deviations from sequence and timing stereotypy did not decrease vocal learning. Moreover, presenting syllables in random sequences enhanced vocal imitation in birds that were tutored later in development. This unexpected finding highlights that sequence complexity can potentiate vocal learning in individuals with more developed vocal control, resembling how complexity and ability interact to modulate various forms of learning in humans.

## INTRODUCTION

Many important behaviors are organized into species-typical temporal patterns. For example, mice and rats engage in stereotyped sequences of grooming behavior that vary across strains and species (Spruijt et al., 1992; Geuther et al., 2021), and different species of songbirds produce vocal sequences with different statistical (sequence) dependencies (e.g., Jin and Kozhevnikov, 2011; Warren et al., 2012; Lipkind et al., 2013; reviewed in Kershenbaum et al., 2014; Cohen et al., 2020; James et al., 2020; Searcy et al., 2022). In addition to the sequences themselves, the timing of elements within behavioral sequences can demonstrate species specificity or typicality (e.g., Fox et al., 2020; Liberman et al., 1958; Araki et al., 2016; Feng et al., 1990; Gerhardt, 2005; Hedwig, 2006; James et al., 2023).

The sensory systems of organisms are tuned to these temporal patterns, with typical sequences and timing leading to stronger behavioral responses or more robust perception compared to atypical patterns (Gerhardt et al., 2007; Comins and Gentner, 2010; Benichov et al., 2016; Taylor et al., 2017; reviewed in Fishbein et al., 2020; Speck et al., 2020). For instance, California thrashers produce particular sequences of song phrases, and males respond more strongly to sequences of phrases that conform to typical patterns than to sequences that violate those patterns (Taylor et al., 2017), and female swamp sparrows prefer acoustic elements (“syllables”) in which the order of constituents (“notes”) conforms to the pattern typical of their local dialect (Balaban, 1988). Male túngara frogs produce a stereotyped sequence of a “whine” followed by a “chuck” (whine-chucks), and the attractiveness of these vocalizations can vary depending on the timing of the whine relative to the chuck (Wilczynski et al., 1999). In addition, increasing or decreasing the silent duration between syllables modulates agonistic responses in indigo buntings (Emlen 1972) and regulates neural responses in various songbird species (Margoliash and Fortune, 1992; Schneider and Woolley, 2013; Lampen et al., 2014; Araki et al., 2016; Bouchard and Brainard, 2016). While there is substantial evidence that temporal patterning affects the strength of behavioral and neural responses, little is known about how temporal patterning affects the acquisition of social and communicative behaviors.

Here we examined the degree to which species-typical temporal patterns regulate vocal learning in the zebra finch. Songbirds like the zebra finch learn their songs during sensitive periods in development (Doupe and Kuhl, 1999; Brainard and Doupe, 2002; Gobes et al., 2019; Sakata and Woolley, 2020), involving both sensory and sensorimotor learning. Zebra finches produce highly stereotyped songs as adults, with acoustic elements (syllables) arranged into stereotyped sequences (motifs) with stereotyped timing (Sossinka and Böhner, 1980; Glaze and Troyer, 2006; Norton and Scharff, 2016; Murphy et al., 2017; James et al., 2023). It has been hypothesized that such stereotypy could facilitate learning because stereotyped sequences of stimuli are more predictable and because more predictable sequences are learned faster by humans (Henson, 1998; Levy, 2008; Smith and Levy, 2013; Clark, 2013; Pearce, 2018; Siegleman et al., 2018; Elazar et al., 2022; Varela et al., 2024). To test whether stereotypy promotes vocal learning in songbirds, we tutored juvenile zebra finches at various ages in development with acoustic stimuli with different degrees of species stereotypy in syllable sequencing or timing.

## RESULTS

We tutored 123 juvenile male zebra finches with stimuli consisting of the same five syllable types using a controlled, operant tutoring paradigm (see Methods; Figure 1a; James and Sakata, 2017; James et al., 2020, 2023). This approach provides a powerful opportunity to reveal factors that modulate the strength of vocal learning. We first analyzed how the age and duration of tutoring modulated the strength of vocal learning and then investigated how the temporal variability of the tutor stimulus interacted with the age at tutoring to regulate learning.

**Figure 1:**
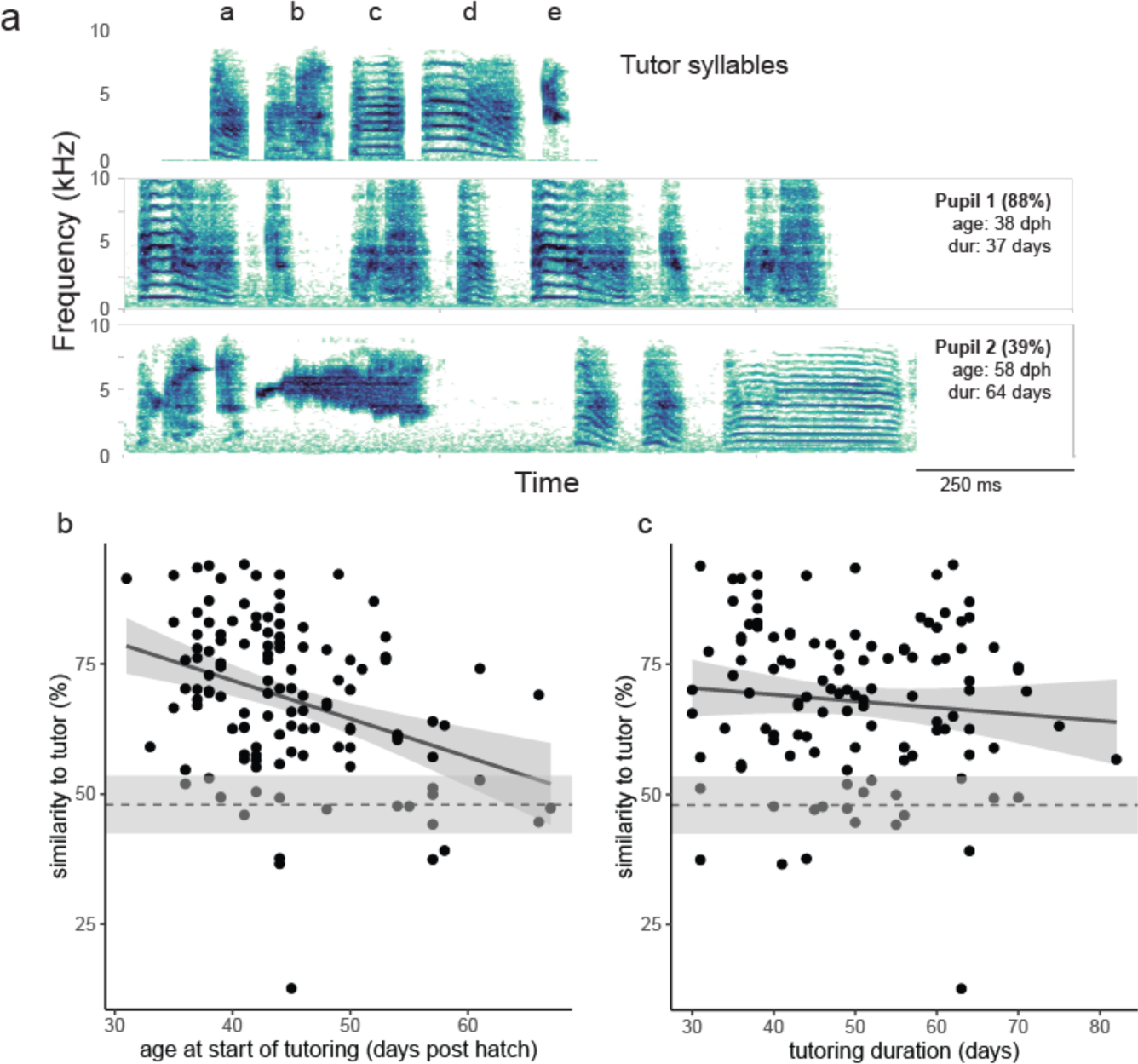
The strength of song learning is affected by the age at which tutoring started but not by the duration of tutoring. (a) Spectrograms of tutor syllables and the adult songs of pupils. Top: spectrogram of the five tutor syllables; middle and bottom: spectrograms of examples of 1-sec snippets of the adult songs of two pupils. The first pupil was tutored early (<50 dph) and produced a song with high similarity to the tutor (88%), while the second bird was tutored later (>50 dph) and produced a song with lower similarity to the tutor (39%). Both birds were tutored with stereotyped (S) stimuli (see Figure 2). (b,c) The age of pupils at the start of tutoring (b; “age at tutoring”) but not the duration of tutoring (c) correlates significantly and negatively with the degree of song learning (i.e., similarity to the tutor song). For both (b) and (c), the dashed horizontal line depicts the mean song similarity [plus 95% confidence interval (CI)] of the songs of untutored birds to the tutor stimulus and represents the chance level of similarity to the tutor song.

### Developmental modulation of song learning

Song is learned during a sensitive period in development that gradually closes over time (Eales, 1985; Brainard and Doupe, 2002; Gobes et al., 2019; Sakata and Woolley, 2020). We tutored juvenile male zebra finches with an identical set of species-typical syllables starting from when they were 31-67 days old (“age at tutoring”). The duration of tutoring ranged from 30-82 days and was statistically independent of the age at tutoring. Therefore, we simultaneously analyzed how age at tutoring and tutoring duration explained variation in the strength of song learning. We computed the acoustic similarity of the adult songs of tutored birds to the tutor stimulus using established software and methods [Sound Analysis Pro (SAP) 2011; “%similarity” as a proxy for the strength of song learning (see Methods)].

Tutored birds demonstrated a wide range of song learning, producing songs that ranged from 13-94% similarity to the tutor stimulus (Figure 1). Age at tutoring (estimate ± error: −0.759 ± 0.174; t_113_=4.35, p<0.0001) but not tutoring duration (−0.140 ± 0.112; t_113_=1.26, p=0.2119) significantly affected the acoustic similarity to the tutor stimulus (multiple regression; Figure 1b,c). Not surprisingly, the older birds were when tutoring began, the less similar their songs were to the tutor stimulus (i.e., inverse relationship between age at tutoring and song learning).

To contextualize the %similarity scores of tutored birds, we computed the acoustic similarity of the adult songs of birds that were not exposed to zebra finch song during the sensitive period for song learning (“untutored birds”; n=20; see Methods) to the tutor stimulus and used these values as a baseline to assess the significance of song learning (i.e., chance level of similarity to the tutor stimulus: e.g., Chaiken et al., 1993; Tchernichovski et al., 2001; Chen et al., 2016; Feher et al., 2017; Chen and Sakata, 2021). Because it is difficult to statistically analyze deviations from baseline values in this manner, we categorized birds based on their age at tutoring, with birds tutored starting from <50 dph classified as “early tutored birds” (n=92) and birds tutored from >50 dph as “late tutored birds” (n=31). [50 dph is approximately halfway between the range of ages of tutoring (31-67).] Both early and late tutored birds produced songs that were significantly more similar to the tutor stimulus than the songs of untutored birds (Dunnett’s test; p<0.01 for each), suggesting significant learning at both developmental periods. However, consistent with the previous analysis, early tutored birds produced songs that were significantly more similar to the tutor stimulus than late tutored birds (t_121_=3.2, p=0.0020).

### General effect of temporal variability of tutor stimuli on song learning

We next analyzed how variability in syllable timing or sequencing affected song learning. The duration of silent gaps between syllables is highly consistent from rendition-to-rendition for adult zebra finch song (e.g., Glaze and Troyer, 2006; Norton and Scharff, 2016; James et al., 2023) and is encoded by neurons in auditory and sensorimotor circuits of songbirds (Margoliash and Fortune, 1992; Lampen et al., 2014; Araki et al., 2016; Bouchard and Brainard, 2016). Zebra finches also produce stereotyped sequences of syllables, and behavioral responses and neural responses within circuits for song learning are sensitive to the sequential order of syllables (e.g., Cynx, 1993; Lewicki and Konishi, 1995; Lewicki and Arthur, 1996; Braaten et al., 2006; Bouchard and Brainard, 2013, 2016; Chen and ten Cate, 2015; Cazala et al., 2019). Given this species-typical stereotypy of timing and sequencing, one group of birds was tutored with stimuli in which syllables were arranged into stereotyped sequences (i.e., same sequence of syllables across all renditions) and with stereotyped gap durations (i.e., all gap durations within the motif and across renditions set to a particular duration)(“stereotyped” or S; n=37). A second group of birds was tutored with a species-atypical stimulus where the gap durations randomly varied from 10-50 ms (in 10-ms increments; with an average of 30-ms gaps)(“variable gaps” or Vgap; n=32), but with stereotyped sequences of syllables. This degree of variability in gap durations across renditions is >5 times greater than the variability observed in typical zebra finch song (James et al., 2023). The third and final group of zebra finches was tutored with a species-atypical stimulus that consisted of variable (randomized) sequences of syllables (“variable sequencing” or Vseq) but with stereotyped gap durations (30-ms gaps; n=54).

Overall, acoustic similarity scores (%similarity) were not significantly different among S-, Vgap-, and Vseq-tutored birds (F_2,120_=1.4, p=0.2603). Compared to untutored birds, S-, Vgap- and Vseq-tutored birds all produced songs that were significantly more similar to the tutor stimulus (F_3,143_=12.2, p<0.0001; Dunnett’s test: p≤0.0001 for each; Figure 2b), indicating significant song learning in response to each type of tutor song organization.

**Figure 2.**
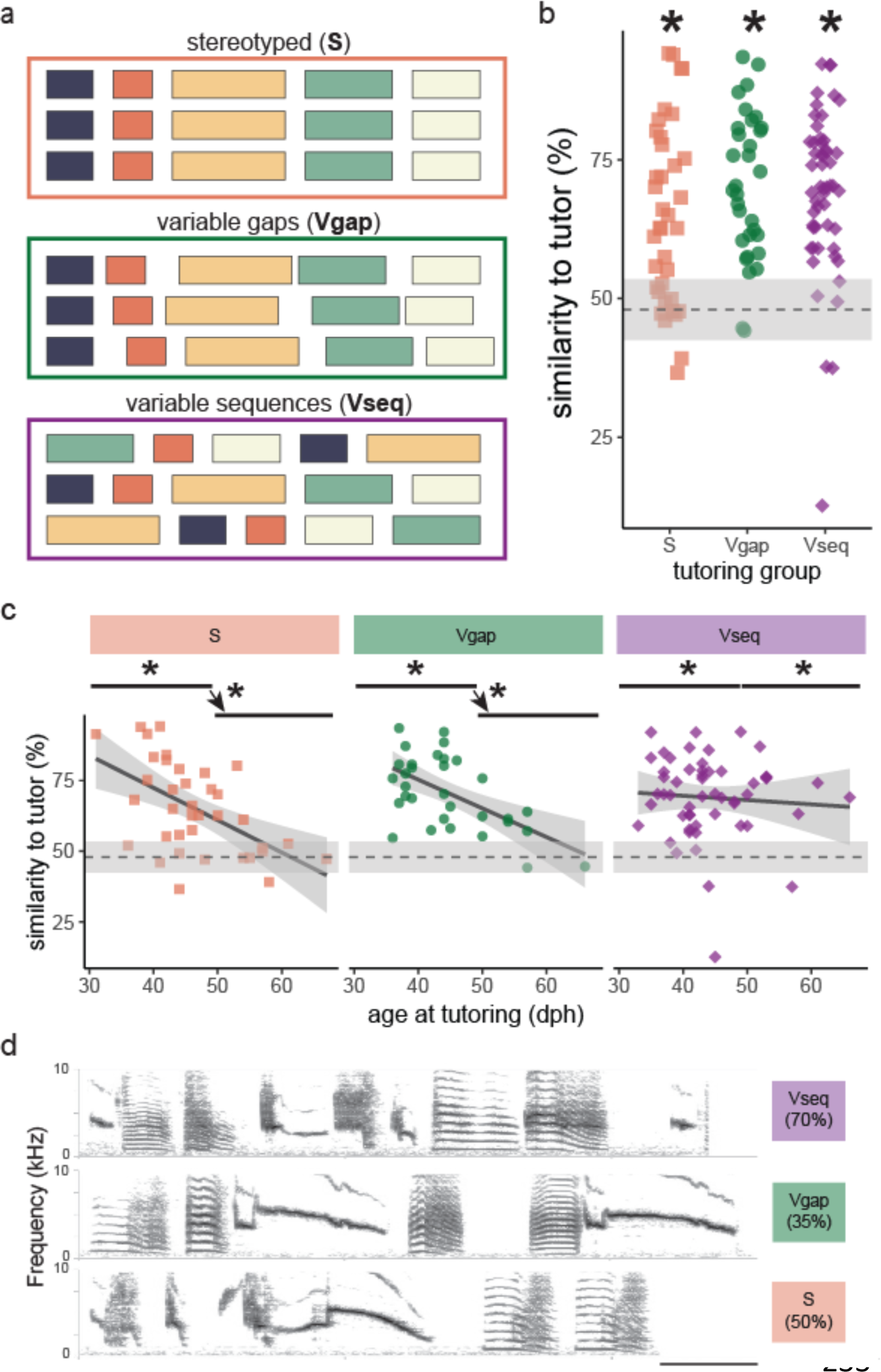
Interactive effects of tutor song organization and age at tutoring on the strength of vocal learning (acoustic similarity to the tutor song). (a) Example schematics of the organization of the three tutor stimuli: stereotyped sequences and gap durations (S), stereotyped sequences with variable gap duration (Vgap), and variable sequences with stereotyped gap durations (Vseq). (b) Tutored birds learn significantly better than chance (i.e., vs. untutored birds), regardless of tutor stimulus organization. Each point depicts the average acoustic similarity of a bird’s song to the tutor stimulus (see Methods). (c) Tutor stimulus organization interacts significantly with the age at tutoring to affect the strength of song learning, with the slope being shallowest in the Vseq group. The bars above each panel denote the age ranges for early tutored birds and late tutored birds. “*” above the bar indicates when distributions are significantly different than untutored birds, and “*” next to arrows indicates when there is a significant difference in similarity to the tutor between early vs. late tutored birds. For (b) and (c), the dashed grey line depicts the mean (and 95% CI) acoustic similarity of the songs of untutored birds to the tutor stimulus (i.e., chance level of similarity). (d) Spectrograms of a single rendition of song from three late-tutored brothers that were tutored with different tutor song organizations. Scale bar= 0.2 sec.

### Interactive effects of temporal variability and age at tutoring on song learning

Given the influence of age at tutoring on the strength of vocal learning (acoustic similarity to the tutor stimulus), we next investigated how the relationship between age at tutoring and vocal learning varied across the three types of tutor song organization. Indeed, it is possible that differences in the efficacy of tutor stimuli are more apparent later in development as learning mechanisms gradually dwindle. Interestingly, there was a significant interaction between tutor stimulus and age at tutoring (ANCOVA: F_2,117_=3.7, p=0.0277; Figure 2c). This appeared to be driven by differences in the strength of the negative relationship among S-, Vgap- and Vseq-tutored birds. When the relationship between song learning and age at tutoring was analyzed within each group individually, the relationship was significantly negative for S- and Vgap-tutored birds (p<0.0010 for each) but not for Vseq-tutored birds (p=0.6073). [Note: there was no significant difference in the age at tutoring among S-, Vgap-, and Vseq-tutoring birds.]

To complement the preceding analyses of the interactive effect of age at tutoring and tutor song organization, we categorically divided tutored birds into early tutored birds (<50 dph) vs. late tutored birds (>50 dph; see above) and examined how song learning varied across tutor song organization and age category (full-factorial model). Consistent with the preceding (ANCOVA) analyses, there was a significant interaction between age category (early vs. late) and tutor song organization (F_2,117_=3.4, p=0.0357). This was driven by the facts that (a) late tutored birds produced songs that were significantly less similar to the tutor stimulus than early tutored birds for S- and Vgap-tutored birds (p<0.01 for each; planned contrast) but not for Vseq-tutored birds and (b) Vseq-tutored birds produced songs that were more similar to the tutor stimulus than S-tutored birds (p=0.0213) or Vgap-tutored birds (p=0.0835) among late tutored birds (planned contrasts) but not among early tutored birds (Figure 2c).

To investigate which groups of birds demonstrated significant learning, we compared the similarity scores of birds tutored early or later in development to the similarity scores of untutored birds in the model. Early tutored birds demonstrated significant learning, regardless of tutor stimulus organization (ANOVA: F_3,108_=14.9, p=0.0001; Dunnett’s: p <0.0001 for each; Figure 2c). However, for late tutored birds, only Vseq-tutored birds demonstrated significant learning (ANOVA: F_3,51_=6.1, p=0.0003; Dunnett’s: p<0.0001; Figure 2c).

One possible reason for the relatively low similarity scores of S- and Vgap-tutored birds that were tutored later in development could be because the songs of S- and Vgap-tutored birds were less mature than Vseq-tutored birds. Less mature songs are more variable from rendition to rendition than more mature songs. Therefore, we analyzed and compared the standard deviation (variability) of similarity scores among late tutored birds. There was no significant difference in song variability among S-, Vgap-, and Vseq-tutored birds (F_2,28_=0.4, p=0.6433), suggesting that songs were comparable in development among late tutored birds with different tutor song organizations.

Song learning and structure have been found to be modulated by genetic factors in various songbirds species, including zebra finches (Forstmeier et al,. 2009; Woodgate et al., 2014; Sato et al., 2016; Mets and Brainard, 2018, 2019; Toji et al., 2023; Shibata et al., 2024). Therefore, we included familyID (father-mother ID; see Methods) as a random factor in the model to control for potential genetic effects on learning. There continued to be a significant interaction between tutor song organization and age at tutoring on song learning (F_2,117_=3.7, p=0.0277), with a significantly negative relationship between song learning and age at tutoring for S- and Vgap-tutored birds (p<0.01 for each) but no relationship in Vseq-tutored birds (p>0.80). Indeed, differences in song learning among late tutored birds tutored with S, Vgap or Vseq stimuli continued to be apparent even after controlling for familyID (see example in Figure 2d).

It was the case that Vseq-tutored birds were tutored for significantly more days than S- or Vgap-tutored birds (F_2,113_=9.1, p=0.0002; Tukey’s HSD: p<0.0200 for each). The preceding analysis of tutored birds (regardless of tutor song organization) suggests no significant influence of tutoring duration on song learning. We extended this previous analysis and analyzed how tutoring duration and tutor stimulus organization (S, Vgap, and Vseq) interacted to affect acoustic similarity to the tutor stimulus (strength of vocal learning) and arrived at similar results that tutoring duration does not influence learning. For example, tutoring duration was not significantly related to vocal learning (estimate ± SEM: −0.13 ± 0.12; t_110_=1.5 p=0.2525), and there was no significant interaction between tutoring duration and tutor stimulus category on vocal learning (F_2,110_=0.1, p=0.9158; i.e., tutoring duration did not affect song learning in S-, Vgap- and Vseq-tutored birds). Other comparisons similarly speak to the lack of effect of tutoring duration on the observed variation in song learning. For example, despite that early tutored birds produced songs that were more similar to the tutor stimulus than late tutored birds, there were no significant differences in the duration of tutoring between early and late tutored birds, regardless of tutoring method (t_114_=0.7; p=0.4742). Relatedly, there was a difference in song learning among S-, Vgap- and Vseq-tutored birds for late tutored birds but not for early tutored birds, but tutoring duration was significantly different among S-, Vgap- and Vseq-tutored birds among early tutored birds (F_2,82_=7.7, p=0.0009) but not late tutored birds (F_2,28_=2.0, p=0.1509). Collectively, these analyses suggest that differences in the duration of tutoring between S-, Vgap-, and Vseq-tutored birds (all were tutored for >30 days) did not contribute to variation in song learning, a finding supported by previous studies (Eales 1985; Peters et al., 1992; Roper and Zann, 2006; Gobes et al., 2019).

### Interactive effects of temporal variability and age at tutoring on syllable learning

Our tutor stimulus consisted of five species-typical syllable “types” that are commonly observed in zebra finch songs (Figure 1a; Zann 1993, 1996): a short, frequency-modulated syllable (“a”), a longer, more complex, and spectrally entropic syllable (“b”; sometimes referred to as a “noise-noise syllable”: Zann 1993), a flat, harmonic stack of intermediate duration (“c”), a longer, complex, frequency-modulated syllable (“d”, a distance call), and a short, high-frequency syllable (“e”). Despite that these species-typical syllables are commonly observed in zebra finch song, it is not clear whether the acquisition and incorporation of these syllables are differentially affected by tutor song organization and age at tutoring. Therefore, we analyzed the degree to which birds tutored with S, Vgap, or Vseq stimuli beginning at different ages produced syllables that resembled each of these tutor syllables (”a”-“e”).

We computed the %similarity score of each pupil’s song rendition to each individual tutor syllable (syllable similarity score) and then averaged these scores over all pupil song renditions. For each rendition, the syllable similarity score reflects the degree to which a syllable similar to the tutor syllable was present in that rendition of a pupil’s song (see Methods). We first ran a full-factorial model with tutor song organization (S, Vgap, and Vseq), age at tutoring, and syllable ID (“a”-“e”) as explanatory variables and found that syllable ID significantly interacted with both tutor song organization and age at tutoring (three-way interaction) as well as just with age at tutoring (p<0.05 for each). Therefore, we next analyzed the interactive effects of tutor song organization and age at tutoring on the %similarity scores for each of the five syllables independently (Figure 3a). For most of the syllables (syllables “a”, “b”, “d”, and “e”), there was a significant effect of age, with syllable similarity scores decreasing as the age of tutoring increased (i.e., pupils were less likely to produce syllables that matched the tutor syllable the later their tutoring started; p<0.05 for each). Furthermore, there was a significant interaction between tutor song organization and age at tutoring for the similarity scores of the “a” syllable (F_2,121_=6.4, p=0.0023), reminiscent of the interaction for song similarity (Figure 2c). For the “a” syllable, this interaction seemed to be driven by a significant negative relationship between syllable similarity scores and age at tutoring for S-tutored birds (p=0.0019) but not for Vgap- or Vseq-tutored birds (p>0.10 for each).

**Figure 3.**
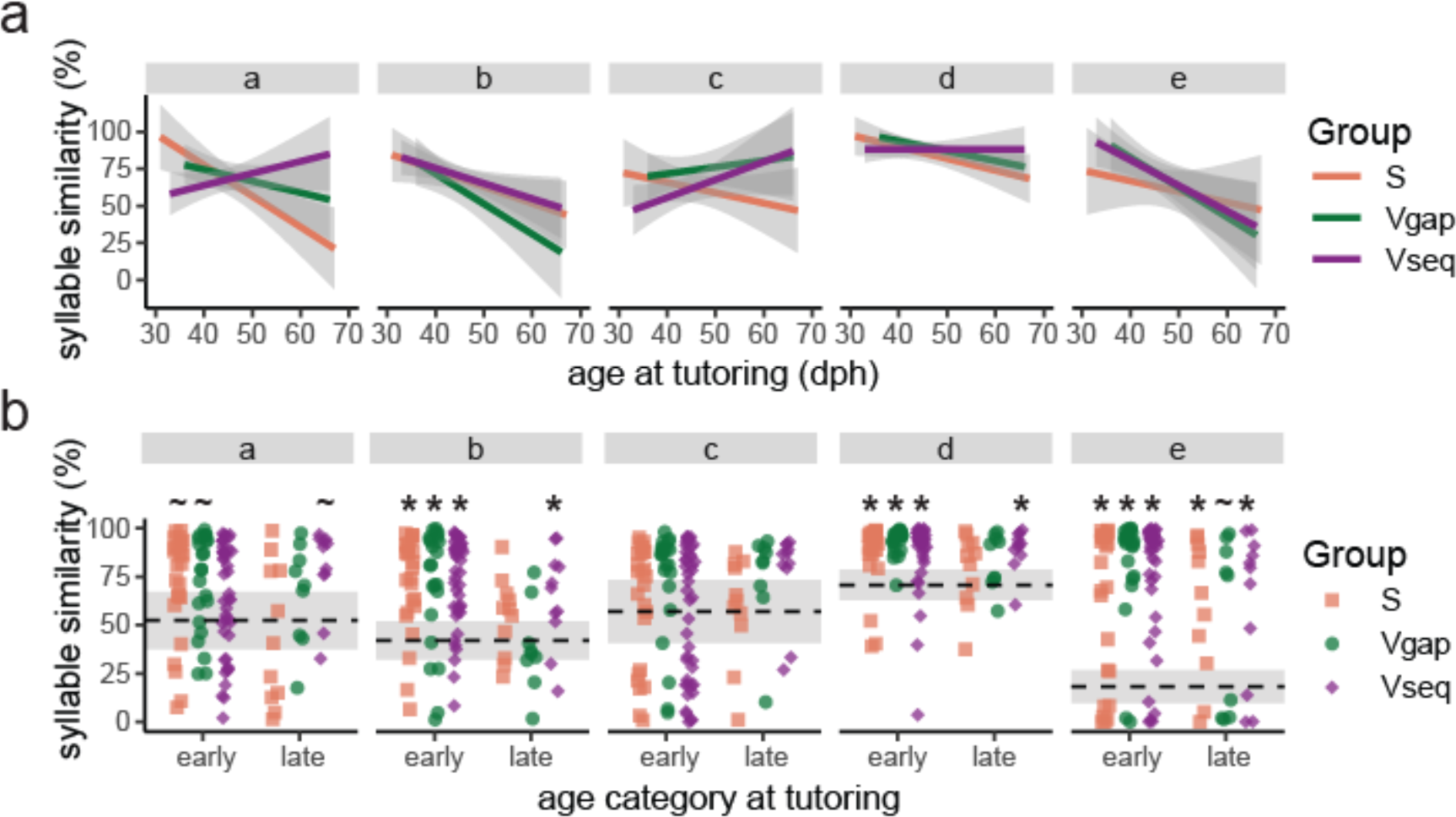
Acquisition of tutor syllables. (a) We analyzed the degree to which syllables within the pupil’s song matched the tutor syllables (“a”-“e”). Specifically, we computed the average %similarity of each tutor syllable to each pupil’s song and correlated this value with the age of tutoring (dph = days post hatch). These relationships were computed for S-, Vgap-, and Vseq-tutored birds independently (see text). Lines plus shaded areas depict the relationship between syllable similarity and age at tutoring. (b). Depiction of raw data for early or late tutored birds according to tutor song organization. The horizontal dashed line represents mean chance level of syllable similarity (i.e., %similarity to each syllable for untutored birds) with the grey area depicting the 95% confidence interval. “*”indicates p<0.05 and “∼”indicates p<0.07 for comparisons to chance (vs. untutored birds).

To gain insight into the significance of syllable learning, we computed the syllable similarity scores for the adult songs of untutored birds to each tutor syllable (i.e., baseline syllable similarity scores), and then compared the syllable similarity scores in S-, Vgap-, or Vseq-tutored birds to these baseline values (see above for similar analysis of song). Among early tutored birds Figure 3b), there were significant differences among the groups for the “b”, “d” and “e” syllables (ANOVA: p<0.05 for each), with S-, Vgap- and Vseq-tutored birds demonstrating significantly higher similarity scores for these syllables (i.e., learning) than untutored birds (Dunnett’s test; p<0.05 for each). There was also a trend for the “a” syllable (p=0.0744), with S- and Vgap-tutored birds tending to have higher similarity scores than untutored birds (p<0.06 for each). Among late tutored birds (Figure 3b), there was evidence of significant differences from baseline for the “b”, “d” and “e” syllables (ANOVA: p<0.05 for each) and a trend for the “a” syllable (ANOVA: p=0.0534). Vseq-tutored birds demonstrated significant learning for the “b”, “d” and “e” syllables and a trend for learning for the “a” syllable (p=0.0693). S-tutored birds demonstrated significantly learning only for the “e” syllable (p<0.01), and there was only a trend of learning of the “e” syllable for Vgap-tutored birds (p=0.0548). In summary, among birds tutored later in development, Vseq-tutored birds demonstrated the most robust evidence of syllable learning.

## DISCUSSION

The sounds that animals use for communication are organized into species-typical patterns. A number of species learn the spectral features of acoustic units for communication (“vocal production learning”), and we tested the hypothesis that deviations away from species-typical temporal patterns would attenuate spectral learning in zebra finches. In addition to previous experiments in songbirds demonstrating such an effect (e.g., Marler and Peters, 1988), we also hypothesized this because the stereotyped nature of syllable sequencing and timing for zebra finch song creates predictability that enhances various forms of learning in humans, including music and speech acquisition (Garrido et al., 2018; Henson, 1998; Levy, 2008; Smith and Levy, 2013; Clark, 2013; Hansen and Pearce, 2014; Bianco et al., 2020; Otsuka and Saiki, 2016; Benitez and Saffran, 2018; Gold et al., 2019; Bianco et al., 2019). Contrary to our expectations, deviations away from species-typical stereotypy in syllable timing or sequencing did not attenuate vocal learning. Moreover, increasing the variability of syllable sequencing (i.e., decreasing stereotypy) enhanced vocal learning: among birds that started tutoring ≥50 dph (late tutored birds), only birds tutored with randomized sequences (Vseq) [and not birds tutored with stereotyped sequences (S) or random gap durations (Vgap)] demonstrated significant song learning. For example, whereas syllable learning for S- and Vgap-tutored birds was primarily significant for birds tutored early in development, Vseq-tutored birds demonstrated significant learning for the “b”, “d” and “e” syllables regardless of age of tutoring. These novel findings not only link the temporal patterning of elements to the fidelity with which individuals learn or imitate those elements (e.g., Marler and Peters, 1988; Plamondon et al., 2008; Mol et al., 2021) but also highlight the role of temporal variability or complexity in engaging mechanisms underlying vocal learning.

Given that sequence variability enhanced the ability of juvenile zebra finches to learn the acoustic structure of syllables, we propose that sequence variability within a tutor stimulus could enhance a juvenile’s attention to the tutor stimulus and, consequently, enhance the sensory encoding of song. Attention has been postulated to play a major role in vocal learning in songbirds as well as humans (ten Cate, 1986; Meltzoff et al., 2009; Chen et al., 2016; Carouso-Peck et al., 2020), and the random sequencing of syllables could prevent habituation to the tutor stimulus, maintain the salience of syllables, and enhance the consolidation of auditory stimuli. Enhancing attention could be particularly important for birds that start to learn songs later in development and could resemble the enhancing effects of social interactions on song learning (Ljubičic et al., 2016; Sakata and Yazaki-Sugiyama, 2020). From a mechanistic perspective, sequence variability can modulate sensory encoding or processing in the sensorimotor nucleus HVC (Bouchard and Brainard, 2016), an area important for the sensory learning of song (Tanaka et al., 2018; reviewed in Ikeda et al., 2020), and it is possible that stimuli consisting of variable sequences lead to patterns of activity or gene expression that promote synaptic plasticity in brain areas underlying song learning.

Birds that were tutored with stimuli with species-atypical levels of temporal variability (Vgap- and Vseq-tutored birds) demonstrated significant song learning. That birds tutored with variable sequences demonstrated significant song learning (i.e., similarity scores significantly greater than chance) is consistent with some previous studies (e.g., James and Sakata, 2017; Lipkind et al., 2017). For example, Lipkind et al. (2017) report that juvenile zebra finches can modify the acoustic structure of their syllables even when syllable sequencing within tutor songs is altered, and James and Sakata (2017) demonstrate that juvenile songbirds tutored with randomized sequences of syllable produce syllables that are classified as accurate imitations of tutor syllables. However, until this study, the degree to which the acoustic structure of the adult songs of birds tutored with variable timing matched the tutor stimulus as well as how various types of temporal variability differentially affected the imitation of syllable structure were not known.

Despite that many studies highlight the importance of predictability to learning, studies also underscore the importance of contextual diversity on speech acquisition and other forms of learning (reviewed in Raviv et al., 2022). The impact of many types of variability (e.g., diversity of training exemplars, of locations and timing of instruction, of contexts, etc.) on perceptual and cognitive processes have been investigated, but one form of variability pertinent to our study is contextual variability. In the domain of word learning, for example, contextual variability can represent the diversity of other words a focal word is produced with. Observations of natural word learning reveal that words emerging first are those that are produced in a wide range of word contexts (Hills et al., 2010; Jones et al., 2012). Further, in experiments of novel word learning, adults are significantly better at recalling and recognizing new words when they appear in a greater diversity of contexts (e.g., in paragraphs on different topics; Johns et al., 2016; Frances et al., 2020). Collectively, these data highlight that contextual variability could not only benefit the acquisition of behaviors as sequentially complex as language but also impact the learning of sequentially simple (stereotyped) behaviors like zebra finch song.

What is particularly interesting about previous findings (and relevant to our findings) is that contextual variability differentially affects learning depending on the age or proficiency of the learner. Low levels of variability (e.g., lower contextual diversity) tend to be more useful in the early stages of learning whereas higher levels of variability can be useful for individuals who have already acquired some proficiency (reviewed in Raviv et al., 2022). For example, in contrast to students with relatively little math training, more experienced math students benefit from exposure to arithmetic problems with high rather than low variability (Likourezos et al., 2019). While it is not the case that variability in sequencing or timing hindered song learning in early tutored juveniles, it is possible that late tutored juveniles benefited from sequence variability in the tutor stimulus because they have already acquired some mastery of vocal control. Juvenile songbirds will practice singing even if they are not provided with the opportunity to hear and memorize songs to imitate, and this practice has been hypothesized to allow juveniles to create an internal model for sound production (Brainard and Doupe, 2013; Mooney, 2022). In this respect, it is possible that older juveniles (>50 dph) are more adept at vocal control and could be in a state to take advantage of the sequence variability in the tutor stimulus.

Our experiments suggest that learning the acoustic properties of syllables might not be significantly affected by variability in gap timing. Variability in gap durations for Vgap-tutored birds was ∼5X as high as that observed in normal zebra finch song, and this degree of variability has been found to affect auditory responses in the zebra finch brain (Lampen et al., 2014, 2019). Despite this variability, Vgap- and S-tutored birds learned the tutor song to a similar degree and displayed similar relationships between song learning and age at tutoring. This lack of effect of gap variability on learning resembles findings from some experiments reporting that auditory perception in zebra finches is less dependent on gap durations than on syllable structure (e.g., Nagel et al., 2010; Dooling and Prior 2017).

In addition to the interactive effects of the temporal variability of tutor song and the age at tutoring on song learning, we also noted variation in the relationship between the age at tutoring and syllable learning across different tutor syllables (“a”-“e”). Although similarity to each tutor syllable generally decreased as the age of tutoring increased for most syllables, the slope was steepest for the “b” and “e” syllables. Zann (1993) classified 2711 elements (from 402 zebra finch males) into 14 different syllable categories or types, and the five syllables used in the tutor stimulus represent the top five categories. These syllable types were found in the songs of 40-90% of the males, with the noise-noise (“b”) and high pitch (“e”) syllable types being the least prevalent of the five. Therefore, the tutor syllable types that were least prevalent in wild birds were the syllable types that showed the steepest diminution in learning with age. This finding is consistent with the experimental finding that zebra finches are more likely to learn common syllable types over uncommon types (ter Haar et al., 2014).

In summary, our findings reveal that the temporal organization of acoustic elements can affect vocal learning in novel ways. In contrast to previous studies describing that syllable timing and sequencing have negligible effects on auditory perception (Lawson et al., 2018; Kriengwatana et al. 2016; Fishbein et al., 2019; Geberzahn and Deregnaucourt, 2020), our expansive study emphasizes that temporal organization can modulate the strength of vocal learning and is consistent with studies highlighting a significant contribution of syllable timing and sequencing to sensory and sensorimotor processes across vertebrate taxa (Bee and Klump, 2005; Verzijden et al., 2007; van Heijningen et al. 2013; Chen and ten Cate 2015, 2017; Benichov et al., 2016; Chen et al. 2016; Spierings and ten Cate 2016; Knowles et al. 2018; Santolin and Saffran, 2018; Rouse et al., 2021; Mol et al., 2021; Ning et al,. 202). Because randomized sequences of tutor syllables seemed to potentiate learning, we propose that sequence variability could prevent neural adaptation, allow for greater attention to acoustic features of syllables, and act upon the same cognitive processes as contextual diversity in humans.

## METHODS

### Animals

Song learning in male zebra finches was analyzed. Birds were provided food and water *ad libitum* and housed on a 14 L:10 D light cycle. All procedures were approved by the McGill University Animal Care and Use Committee in accordance with the guidelines of the Canadian Council on Animal Care.

Birds were raised by both parents in a sound-attenuating chamber (“soundbox”; TRA Acoustics, Ontario, Canada) up to 5-7 days of age, at which time their father was removed. Only male zebra finches learn and produce courtship song, and the sensitive period for song learning starts at approximately 20 days post-hatching (Brainard and Doupe, 2013; Roper and Zann, 2006). Thus, juveniles were not exposed to song from a live adult during the sensitive period for song learning and were naïve to song before experimental tutoring. When these juveniles were nutritionally independent (i.e., could feed themselves; ∼30–40 days old), they were housed individually in a soundbox for a period of song tutoring (see below) and thereafter until they were mature (∼4 months of age; Derégnaucourt et al., 2005; James and Sakata, 2017; Lipkind et al., 2013; Lipkind et al., 2017; Mets and Brainard, 2018; Mets and Brainard, 2019; Tchernichovski et al., 2001; James et al., 2023).

### Song tutoring and stimuli

Birds were operantly tutored for >1 month with song playback operantly triggered by perch hops (n = 121) or string pulls (n = 2) using custom-built perches or strings connected to a National Instruments PCI-6503 I/O card (National Instruments, TX). Each hop or string pull triggered the playback of one song (i.e., four concatenated motifs of the tutor stimulus; see below). Song playbacks were spaced in time such that juveniles could hear a maximum of 10 operantly triggered playbacks within each of three time periods per day (morning, noon, and afternoon); this schedule of song exposure has been found to enhance song learning (Tchernichovski et al. 1999; Derégnaucourt et al. 2005). We used Sound Analysis Pro 2011 (SAP; http://soundanalysispro.com) for song tutoring, and stimuli were played out of an Avantone Pro Mixcube speaker (Avantone, NY) connected to a Crown XLS 1000 amplifier (Crown Audio, IN).

Birds were between 31-67 dph when tutoring started (“age at tutoring”). Most of the birds were tutored between 30-50 dph because birds learn best during this period and because many of these birds were tutored for experiments that investigated common song patterns in learned song (i.e., we aimed to maximize learning: James and Sakata, 2017; James et al., 2020, 2023). Importantly, the age at tutoring was not significantly different among S-, Vgap-, and Vseq-tutored birds (F_2,120_=1.2, p=0.2967).

Birds were tutored, on average, for 49.7 ± 11.5 days (mean ± S.D.; range: 30 - 82). The duration of tutoring was unavailable for seven birds (i.e., the start of tutoring was noted but not the end of tutoring). All of these birds were Vseq-tutored birds that started their tutoring between 35 - 42 dph (i.e., early tutored birds), and their similarity scores were well within the range of Vseq-tutored birds with information on tutoring duration (range: 55-90%). Given the large sample size of the current analysis correlating song learning with tutoring duration (n=116 birds) and that previous studies similarly document a lack of effect of tutoring duration on learning for birds tutored for at least a week (Eales, 1985; Roper and Zann, 2006; Gobes et al., 2019), we do not anticipate the absence of data from these birds will affect the observed relationships between the ages and durations of tutoring and similarity scores in the existing dataset. For example, when the data for the seven birds without information on tutoring duration were removed, the interactive effects between the age at tutoring and tutor song organization on song (snippet) learning (F_2,110_=3.5, p=0.0333) and between age at tutoring and syllable ID on syllable learning (F_4,440_=4.7, p=0.0011) remained significant, as did patterns in syllable learning for early and latetutored birds; therefore, these birds remained in the dataset for the reported analyses.

All song stimuli consisted of five canonical zebra finch song elements (“syllables“; hereafter, “a”, “b”, “c”, “d”, and “e“; Figure 1). Zebra finch song is typically organized into bouts (seconds in duration) that contain multiple renditions of a motif (stereotyped sequence of syllables that are, on average, 0.6-0.7 sec: Sossinka and Bohner, 1980; Zann, 1993, 1996; Tchernichovski et al., 1999; Franz and Goller, 2003; Whitney and Johnson, 2005; Glaze and Troyer, 2006, 2013; Pytte et al., 2007; Holveck et al., 2008; Funabiki and Funabiki, 2009). Song stimuli consisted of four renditions of a motif to emulate species-typical song bouts. In addition, because gaps are longer between motifs than within motifs (Glaze and Troyer 2006; Hyland Bruno and Tchernichovski 2019), between-motif gap durations were fixed at 100 ms for all stimuli. The variability of syllable sequencing and timing across motifs distinguished the experimental groups.

Birds that were exposed to stereotyped (S) stimuli were tutored with stimuli in which the sequence of syllables and duration of silent gaps between syllables within the motif were fixed across all renditions of song (n=37). For instance, a bird might be tutored solely with the sequence “becad” with 30-ms gaps between each syllable within the motif (see Supplementary Table 1). Ten different motif sequences were used to tutor birds in the S group, and inter-syllable gaps were fixed at 10, 20, 30, 40, or 50 ms (with each bird being tutored with only one of these sequences and generally a single inter-syllable interval: Supplementary Table 1). These durations were chosen to cover the species-typical range of gaps observed in zebra finches (Price 1979; Norton and Scharff 2016; Cooper and Goller 2006; Norton and Scharff 2016; Lachlan et al. 2016; Araki et al. 2016; James and Sakata 2017). Three S-tutored birds were tutored with gap durations that did not vary across renditions of the motif but varied between different syllables in the motif; specifically, two birds were tutored with the sequence “becad” and one bird with the sequence “dacbe” in which the four gaps between syllables within the motif were fixed, respectively, at 20, 50, 10, and 40 (Supplementary Table 1).

Another group of birds was tutored with stereotyped sequences of syllables in which the duration of the gaps varied randomly within motifs and across renditions of motifs (“variable gap” or Vgap stimulus; n=32). We manipulated gap durations because neurons in auditory and sensorimotor circuits of songbirds are sensitive to the duration of silent gaps between syllables (Margoliash and Fortune, 1992; Lampen et al., 2014; Araki et al., 2016; Bouchard and Brainard, 2016) and manipulated the variability of gap durations because gap durations are typically highly consistent from rendition-to-rendition for adult zebra finch song (Glaze and Troyer, 2006; Norton and Scharff, 2016; James et al., 2023); for example, across 43 adult zebra finches, median gap durations were ∼40 ms and the median IQR for these gaps was 3-4 ms: L.S.J. & J.T.S, unpublished data). Each bird was presented with a single sequence of syllables (e.g., “becad”), but the gap durations between syllables within the motif were randomly selected from 5 possible gap durations: 10, 20, 30, 40, or 50 ms. Zebra finches attend to, discriminate among, and imitate gap durations in this range of gap durations (e.g., James et al., 2023). Gap durations were pseudo-randomly chosen such that each gap duration was presented the same number of times; as such, the average gap duration was 30 ms for these stimuli. Twelve different motif sequences were used to tutor birds in the Vgap group (Supplementary Table 1), only two of which differed from those used as stimuli for S-tutored birds.

The last group of birds were tutored with stimuli in which the sequence of the syllables within the motif varied from rendition to rendition (including within the same song bout; “variable sequencing” or Vseq stimulus; n=54). Specifically, we first synthesized 120 different 5-syllable sequence variants in which each syllable is produced only once. Sequences were created in this manner because zebra finches do not typically repeat syllables within the motif; therefore, sequences in which each syllable is produced only once is typical of this species. We then pseudo-randomly selected and concatenated four different sequence variants to construct a song bout, ensuring that each of the 120 sequence variants was presented equally. Gap durations within the motif were fixed at 30 ms for all birds in the Vseq group; this was the average gap duration within the stimuli of Vgap-tutored birds and the most common gap duration for the stimuli of S-tutored birds.

In addition to tutored birds, some birds were raised without exposure to conspecific song (“untutored birds”; n=20) during the sensitive period for song learning; they were raised with their mother and then housed individually until their songs were recorded in adulthood (see below). These birds served to compute chance levels of similarity to the tutor stimuli (i.e., to assess the significance of song learning: e.g., Chaiken et al., 1993; Tchernichovski et al., 2001; Chen et al., 2016; Feher et al., 2017; Chen and Sakata, 2021;).

### Song recording and analysis

When tutored birds were sexually mature (mean ± S.D.: 121.6 ± 8.7 dph; range: 87 - 139), they were individually housed in a soundbox for song recording. Age at song recording was not significantly different among the three groups of tutored birds (F_2,120_=0.7, p=0.9301), or related to the mean or variability of %similarity scores of tutored birds (p>0.25 for each). Untutored birds were slightly older at the time of song recording (mean ± S.D.: 132.0 ± 20.2 dph; range: 92 - 164); this allowed for more time for their songs to stabilize.

For all song recordings, birds were housed individually in a soundbox, and song was recorded using an omnidirectional microphone (Countryman Associates, Inc, Menlo Park, CA) positioned above the bird’s cage. A computerized, song-activated recording system was used to detect and digitize song (Sound Analysis Pro (SAP) 2011; digitized at 44.1 kHz). Recorded songs were digitally filtered (0.3–10 kHz) for off-line analysis using software custom-written in the Matlab programming language (MathWorks, Natick, MA). All songs recorded and analyzed were spontaneous songs produced in isolation (“undirected song”).

After identifying recordings with songs, we annotated introductory notes and song syllables within song bouts. Zebra finches repeat a series of short, acoustically simple syllables called introductory notes before singing their song. Even untutored birds and birds tutored without introductory notes in the stimulus produce readily identifiable introductory notes in their songs (e.g., Price, 1979; Kalra et al., 2021). Many studies quantify song learning by first identifying a stereotyped sequence of syllables (motifs) in pupils. However, a number of pupils in this study produced adult songs with variable acoustic and temporal structure (i.e., some birds did not produce a stereotyped motif at the time of their adult song recording); this made motif identification difficult and subjective. Consequently, we extracted the first 1-sec of song (excluding the repetition of introductory notes) ensuring that we retained the entire syllable that first crossed the one second mark (i.e., we did not clip any syllables). Zebra finch motifs are, on average, 0.6-0.7 sec (Sossinka and Bohner, 1980; Zann, 1993, 1996; Tchernichovski et al., 1999; Franz and Goller, 2003; Whitney and Johnson, 2005; Glaze and Troyer, 2006, 2013; Pytte et al., 2007; Holveck et al., 2008; Funabiki and Funabiki, 2009). Therefore, a one second snippet is ∼50% longer than the average motif and is likely to contain a full motif. We compared the acoustic similarity of syllables in this snippet to the tutor syllables, providing a relatively unbiased approach to examine song learning across birds, regardless of the stereotypy of their adult song. The focus on song structure following introductory notes is useful because tutor stimuli were not preceded by the repetition of introductory notes.

We evaluated the acoustic similarity between song snippets and a motif of the tutor stimulus using SAP. Specifically, we computed the %similarity score (using the “asymmetric” and “time course” settings) for each song snippet of a pupil, with a tutor motif as a reference. This measurement uses acoustic features (pitch, frequency modulation, amplitude modulation, goodness of pitch, and Wiener entropy) to compute how much of the tutor stimulus is present in the pupil’s song. Importantly, this similarity metric is useful to quantify song learning here because it is not affected by the sequence of syllables in the tutor stimulus or by multiple examples of a syllable within the song snippet and does not factor in silent gaps in the calculation. We used the default settings on SAP that are optimized for zebra finch song (p=0.05, interval=70 ms; minimum duration=15 ms). We measured the SAP similarity for, on average, 36.7 ± 7.9 song snippets per pupil (mean ± S.D.; range: 27 - 77), and computed the mean %similarity score across all song snippets for each pupil for analyses. Sample sizes were similar for untutored birds (mean + S.D.: 31.9 ± 10.4; range: 12 – 51). These ranges of sample sizes are comparable to many other studies of song learning. The number of snippets was not significantly related to the mean or variability of %similarity scores for tutored birds (p>0.3 for each). To identify variation in the learning of each of the five individual tutor syllables, we ran the same similarity measurements across all song snippets except with the tutor stimulus limited to a single syllable (e.g., “a”, “b”, “c”, etc).

Aspects of the sequencing, timing, and acoustic structure of the songs of many of these birds have been published previously (James and Sakata, 2017; James et al., 2020, 2023). In particular, convergence in syllable sequencing (i.e., learning bias) was documented in Vseq-tutored birds in James and Sakata (2017); the contribution of motor production biases to syllable sequencing in Vseq-tutored and 2 untutored birds were described in James et al. (2020); and fidelity and biases in the learning of gap durations was documented in James et al. (2023). The acoustic similarity of the motifs of a subset of the Vseq-tutored birds (n=45 out of 54) to the tutor stimulus have been reported in the Methods section of James and Sakata (2017) to quantify the efficacy of our tutoring and to compare the level of learning to other studies, and the SAP similarity scores of motifs of some S-tutored birds (n=13) that produced a stereotyped motif were reported in Kalra et al. (2021).

Given that some aspects of the songs of many of the birds in this study were reported in previous studies, it is important to discuss the novelty of the current analyses. In none of the previous reports was the similarity of song snippets or of individual syllables to the tutor syllables explicitly analyzed. The acoustic similarity of the songs of Vgap- and most of the S-tutored birds to the tutor stimulus have not been examined before, and in no previous study has the similarity scores been compared across the different tutoring regimes or across age at tutoring. Further, data from 8, 4, and 5 additional birds were added to the S-, Vgap-, and Vseq-tutored groups, respectively, for the current analysis; these birds were tutored after the completion of the aforementioned studies to increase sample sizes.

### Statistical analyses

In general, we used ANOVAs to analyze the effects of tutor stimulus (S, Vgap, and Vseq) on song learning. We also used regressions, multiple regressions, and ANCOVAs to analyze how age at tutoring and duration of tutoring independently affected song learning or interacted with tutor song organization to affect song learning. Analyses were also run with pairID (a unique code for male-female pairings; to control for potential genetic contributions) as a random factor. To compare between groups of tutored birds, we used Tukey’s HSD tests, and to compare between untutored birds and each group of tutored birds, we used Dunnett’s tests. All analyses were run in JMP v.17 for the Mac, with α=0.05 for all analyses. Figures were generated in R (v.4.2.1) using RStudio v.2023.12.1.

## Acknowledgements

We would like to thank F. Bendall, V.Y. Li, and A.S. Wang for their assistance with data collection and processing, and M.S. Brainard and S.C. Woolley for comments on manuscripts and data analysis and interpretation. This work was supported by funding from the National Science and Engineering Research Council (Discovery Grant #05016 to J.T.S.; USRA fellowship to O.R.), the Fonds de Recherche du Quebec – Nature et technologies (PR-299652 to S.C.W.), Smithsonian Institute Postdoctoral Fellowship (L.S.J.), and the Centre for Research on Brain, Language and Music (L.S.J. and J.T.S).

