## Supplementary Table 1 for "Temporal variability enhances the acquisition of stereotyped communication signals"

**Supplementary Table 1.** Number of birds tutored with specific syllable sequences (motifs) and gap durations between each syllable in the motif.

| Sequence | 10 | 20 | 30 | 40 | 50 | Different gaps between syllables | Variable gaps across renditions |
|---|---|---|---|---|---|---|---|
| **variable** | | | n = 54 | | | | |
| **abced** | | | | | | | n = 1 |
| **abdce** | | | | | | | n = 1 |
| **abdec** | | | n = 1 | | | | n = 3 |
| **aecbd** | | | n = 2 | | | | n = 1 |
| **bacde** | | n = 1 | n = 2 | | n = 1 | | n = 3 |
| **beadc** | n = 1 | | n = 4 | | n = 1 | | n = 4 |
| **becad** | | n = 1 | n = 1 | | n = 1 | n = 2 | n = 4 |
| **cdaeb** | n = 1 | | n = 2 | n = 1 | n = 2 | | n = 4 |
| **cedba** | | | | | | | n = 1 |
| **dacbe** | n = 2 | n = 1 | n = 1 | n = 1 | n = 1 | n = 1 | n = 5 |
| **decab** | | | n = 2 | | | | |
| **eabcd** | | | n = 1 | | | | n = 2 |
| **edbca** | | | n = 1 | | n = 2 | | n = 3 |

*Note:* The gap duration of 30 ms is overrepresented in the variable sequence group because this was the average gap duration between syllables for the VG-tutored, and the most common gap duration for the stimuli of S-tutored birds. The sequences “becad” and “dacbe” which the four gaps between syllables within the motif were fixed, respectively, at 20, 50, 10, and 40 most resemble typical zebra finch song structure.
